# Population effect of influenza vaccination under co-circulation of non-vaccine variants and the case for a multi-strain A/H3N2 vaccine component

**DOI:** 10.1101/054403

**Authors:** Colin J Worby, Jacco Wallinga, Marc Lipsitch, Edward Goldstein

**Affiliations:** Department of Epidemiology, Center for Communicable Disease Dynamics, Harvard TH Chan School of Public Health, Boston MA, 02115 USA; National Institute of Public Health and the Environment (RIVM), Antonie van Leeuwenhoeklaan 9, 3721 MA Bilthoven, The Netherlands; Leiden University Medical Center, Department of Medical Statistics and Bioinformatics, 2300 RC Leiden, Netherlands; Department of Immunology and Infectious Disease, Harvard TH Chan School of Public Health, Boston MA, 02115 USA

**Keywords:** Influenza, co-circulating strains, monovalent vaccine, bivalent vaccine, cross-immunity

## Abstract

Some past epidemics of different influenza (sub)types (particularly A/H3N2) in the US saw co-circulation of vaccine-type and variant strains. There is evidence that natural infection with one influenza (sub)type offers short-term protection against infection with another influenza (sub)type (henceforth, cross-immunity). This suggests that such cross-immunity for strains within a (sub)type is expected to be strong. Therefore, while vaccination effective against one strain may reduce transmission of that strain, this may also lead to a reduction of the ability of the vaccine-type strain to suppress spread of a variant strain. It remains unclear what the joint effect of vaccination and cross-immunity is for co-circulating influenza strains, and what is the potential benefit of a bivalent vaccine that protects against both strains.

We simulated co-circulation of vaccine-type and variant strains under a variety of scenarios. In each scenario, we considered the case when the vaccine efficacy against the variant strain is lower than the efficacy against the vaccine-type strain (monovalent vaccine), as well the case when vaccine is equally efficacious against both strains (bivalent vaccine).

Administration of a bivalent vaccine results in a significant reduction in the overall incidence of infection compared to administration of a monovalent vaccine, even with lower coverage by the bivalent vaccine. Additionally, we found that the stronger is the degree of cross-immunity, the less beneficial is the increase in coverage levels for the monovalent vaccine, and the more beneficial is the introduction of the bivalent vaccine.

Our work exhibits the limitations of influenza vaccines that have low efficacy against non-vaccine strains, and demonstrates the benefits of vaccines that offer good protection against multiple influenza strains. The results elucidate the need for guarding against the potential co-circulation of non-vaccine strains for an influenza (sub)type, at least during select seasons, possibly through inclusion of multiple strains within a (sub)type (particularly A/H3N2) in a vaccine.

## Introduction

The recurrence of seasonal influenza epidemics is driven by a number of factors including waning of immunity, weather-related changes in transmissibility of influenza [1], and antigenic changes in the influenza virus [2]. Antigenic change creates a need for an update of influenza vaccines for each hemisphere every year to several years [3]. Despite those updates, there is significant circulation in some years of influenza strains for which the vaccine offers little protection. The most recent instance of such circulation in the US and elsewhere, took place during the 2014-15 influenza season. During that season, vaccine-type A/H3N2 viruses (that is, A/Texas/50/2012-like viruses) were a majority of A/H3N2 isolated in the US early in the season (up to week 47) [4]; by week 50, the vaccine-type strain had declined to about 30% of A/H3N2 specimens [5], with the remainder either showing reduced titers to vaccine-derived antisera or belonging to a genetic lineage showing such reduced titers. The decline in the proportion vaccine-type among A/H3N2 continued through the rest of the season [6]. This predominance of the novel A/H3N2 strain also contributed to the very low vaccine effectiveness against influenza A/H3N2 during the 2014-2015 season [7]. The overall effectiveness of vaccination against influenza A/H3N2 during 2014-5 in the US was unusually low at 13%; this overall low effectiveness was a combination of 43% effectiveness against vaccine-like virus and 9% effectiveness against “vaccine-low” viruses [8].

Competition between co-circulating strains within an influenza (sub)type was observed during previous seasons as well, with a variety of outcomes. During the 2004–05 season, vaccine-type viruses initially dominated the 2004–05 A/H3N2 incidence [9], but they were subsequently replaced by a non-vaccine strain [10]. Vaccine effectiveness during that season was very low [7]. When the Fujian A/H3N2 strain appeared in the US during the 2003–2004 season, that strain had dominated the circulation of influenza, and the proportion of the vaccine strain declined during the course of that season (compare [11] to [12] to [13]), though the decline in the proportion of the vaccine-type strain among the A/H3N2 specimens (from 25% [11] to 11.5% [13]), was not as drastic as for the 2004-05 season. Moreover, while vaccine effectiveness during the 2003–04 season was relatively low, it was somewhat higher than during 2004–05 season [14]. During the 2007–2008 A/H3N2 season, an antigenic variant of the vaccine strain circulated at higher levels compared to the vaccine-type strain [15]. However, the relative share of those two strains varied little through the course of the season [16]. During the 2011–2012 influenza B season, over half the detected B specimens were not of the vaccine type [17]. The proportion of vaccine and non-vaccine-type viruses did not seem to change through the course of that season [18].

When a non-vaccine strain co-circulates and vaccine effectiveness against it is low (as in 2004–5 and 2014–5), it is commonly thought that vaccination reduces the incidence of the vaccine-type strain and has limited impact on mitigating the incidence of a non-vaccine strain. An additional effect of vaccination that is often neglected is a potential increase in the incidence of a non-vaccine strain through reduction of the incidence of the vaccine-type strain, cutting down on the mitigating effect of the incidence of vaccine-type strain on the incidence of the non-vaccine strain through cross-immunity. This cross-immunity, which translates into the reduction in the risk of infection with one influenza strain for a period of time following an infection with another influenza strain, is believed to be conferred by a variety of immunological mechanisms, and its consequences are documented in the literature. Sonoguchi et al. [19] studied the impact of the same-season circulation of A/H3N2 and A/H1N1 influenza in Japanese schools, concluding that infection with A/H3N2 was negatively associated with subsequent risk of infection with A/H1N1. Cowling et al. [20] have found that those infected with seasonal influenza A during the 2008–2009 season in Hong Kong had a lower risk of laboratory-confirmed pandemic A/H1N1 infection. The results in [20] were further extended to show shortterm cross-protection against infection by unrelated viruses [21]. Ferguson et al. [22] and Tria et al. [23] concluded that strong, transient, nonspecific immunity effective against all influenza strains was necessary to produce realistic patterns of sequence diversity in simulations of influenza A and B evolution. Epidemiological consequences of cross-immunity between different influenza (sub)types were demonstrated in [24]. That paper has shown that the magnitude of early population incidence of some influenza (sub)types is negatively correlated with the cumulative seasonal incidence of other influenza (sub)types. While we are unaware of any studies directly addressing cross-immunity within a season and within a subtype, it is expected it to be even stronger than cross-immunity for different influenza (sub)types, rendering infection twice in the same season with the same subtype quite unlikely. Though untested, this hypothesis seems plausible in light of the strong evidence for cross-immunity between influenza (sub)types.

Given cross-immunity within a subtype, the impact of vaccination when there is cocirculation of a non-vaccine strain is uncertain, and the dependence of that impact on the strength of cross-immunity is unclear. In this paper, we explore these issues by simulations of influenza transmission in an age-stratified population under a variety of scenarios for transmission parameters and vaccination coverage levels. While some of the choices we make are motivated by data from recent epidemics in the US, the aim of this work is not to calibrate transmission models to the actual epidemic data but rather to establish general principles of the interaction of vaccination and cross-immunity for co-circulating influenza strains. The ultimate goal is the elucidation of the need to guard against the co-circulation of non-vaccine strains (particularly for influenza A/H3N2) for which vaccine efficacy is low, possibly by employing bivalent vaccines. In fact, a precedent for this exists, as continuing co-circulation of the Victoria (vaccine-type) and the Yamagata influenza B lineages led to the introduction of a quadrivalent influenza vaccine containing both strains starting with the 2013–2014 season, though no bivalent A/H3N2 vaccine component has ever been adopted by the WHO.

## Materials and methods

*Outline:* We simulate influenza outbreaks in an age-stratified population for two co-circulating strains (which may be introduced to the population at different times). We compare the performance of several different policies defined by the vaccine coverage and the valency of vaccine used – these are the two variables that are under the control of a policy maker (Table 1). A *monovalent* vaccine has a lower efficacy against one of the two strains than the other, while a *bivalent* vaccine has equal efficacy against both strains. The outcome considered in the policy comparison is cumulative incidence of infection over the course of a season. The policy comparison is made across a set of scenarios -each scenario defined by one combination of values for the parameters not under a policy maker’s control: the degree of cross-immunity conferred by natural infection with one strain against subsequent infection with another, and several properties that affect the transmission dynamics of the two strains (Table 2). Each policy is compared against a *baseline* policy of using the monovalent vaccine with 40% coverage for children, 30% for adults, similar to recent US data. We examine the scenarios when the monovalent vaccine administration varies (is either reduced or increased relative to the baseline levels, see Table 1) and compare them to the baseline. Additionally, we compare the baseline scenario with a monovalent vaccine to scenarios in which a *bivalent* vaccine (with equal efficacy against both strains) is used, the same range of coverage levels as for the monovalent vaccine (Table 1), again comparing outbreak size with the baseline coverage of monovalent vaccine. We report the comparisons separately for three different values of the cross-immunity parameter θ, which, as defined in the Transmission Model subsection, is the degree of cross-protection offered by natural infection.

**Table 1:** Coverage levels considered for both the monovalent and the bivalent vaccines in our simulations.

| Coverage scenario | Monovalent vaccine coverage | Bivalent vaccine coverage |
| --- | --- | --- |
| Uniform coverage | 100% children/100% adults | 100% children/100% adults |
| 50% increase | 60% children/45% adults | 60% children/45% adults |
| 25% increase | 50% children/37.5% adults | 50% children/37.5% adults |
| 10% increase | 44% children/33% adults | 44% children/33% adults |
| Baseline level | 40% children/30% adults<br>(*Baseline case*) | 40% children/30% adults |
| 10% reduction | 36% children/27% adults | 36% children/27% adults |
| 25% reduction | 30% children/22.5% adults | 30% children/22.5% adults |
| 40% reduction | 24% children/18% adults | 24% children/18% adults |

We define the following parameters – vaccine coverage levels, valency of vaccine and cross immunity -to be ‘primary parameters’ (T_1_), and all remaining parameters governing transmission dynamics ‘secondary parameters’ (T_2_). In order to perform the comparisons described in Table 1, we repeatedly draw plausible values of the secondary parameters, T_2_, based on estimates form the literature (Table 2, rows 4–10), to generate a large collection *S* of those parameters. For each set of secondary parameters in the collection *S*, we calculate the cumulative incidence of infection over the course of a season for each of the sixteen vaccine policies described in Table 1, and for three levels of cross-immunity θ, giving a total of 48 sets of the values of the primary parameters. We note that with a deterministic model as used here, a choice of the primary and secondary parameters T_1_ and T_2_ completely defines an epidemic in the community. For each level of cross immunity, the outcome under each of fifteen alternative vaccination policies (described in Table 1) are compared to the outcome with baseline coverage of monovalent vaccine.

*Transmission model:* We use the notation described in the Outline sub-section. We consider two strains, 1 and 2, with 1 being the target of the monovalent vaccine, which has lower efficacy against 2. We use a deterministic, difference equation model with a daily time step, modeling the spread of these strains in an age-stratified population with 5 age groups (0-4,5-17,18-49,50-64,65+). Transmission dynamics are modeled in the stratified mass action two-strain SIR (S,I1,I2,R) framework [25] (with the parameters described in Tables 2 and 3). Contacts between the different age groups (strata) are described by a symmetric matrix *C* = (*c_ij_*), where *c_ij_* is the average number of contacts per unit of time (day) between a pair of individuals in strata *i* and *j*. We estimate the matrix *C* using the POLYMOD study data [26,27]. Additionally, for each age group *i*, we have

A. Population size (extracted from [28], based on the 2014 US population).
B. Individual relative susceptibility 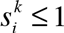 (per contact with an infected individual) for strain *k* for each individual in stratum *i* (uniform susceptibility). We assume that for *i* ≤ 3, 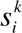 is drawn uniformly between [0.75,1]; 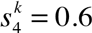; 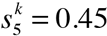

Based on previous work [29], we assume that infectivity is age-independent. We assume that an infector causes infections in the community for 7 days following his/her own infection, and the distribution *ω()* of infectiousness over those 7 days (serial interval distribution) is borrowed from [29]. Thus, in the absence of vaccination, the number of infections during the early stage of an epidemic in age group *i* caused by a person in age group *j* on day *d* (1 ≤ *d* ≤ 7) after that person’s own infection is

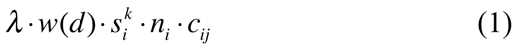

The leading eigenvalue of the next generation matrix 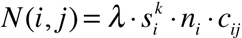 is the initial effective reproductive number in the absence of vaccination. We fix λ so when 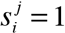 for *i* ≤ 3 (maximal possible susceptibility), the initial effective reproductive number (in the absence of vaccination) is 1.4 for both the vaccine and the non-vaccine strains.

Based on data for vaccination levels before the start of seasonal epidemics in recent years in the US, we assume that in the baseline case 40% of children and 30% of adults are vaccinated. Little is known about the efficacy of influenza vaccine against infection. The annual estimates published by the US CDC refer to effectiveness against symptomatic, PCR-confirmed infection episodes. Our previous work [30] has suggested that efficacy of some vaccines against infection can be lower than efficacy against symptomatic disease. We assume that vaccine reduces susceptibility to a vaccine strain by 40% for non-elderly (age groups 1–4) and by 30% for the elderly. For the “monovalent” vaccine type, we assume that susceptibility to non-vaccine strain is reduced by *x_1_*% for non-elderly, *x_2_*% for elderly, where *x_1_* is drawn uniformly between [0,20] and x_2_ drawn uniformly between [0,15].

We assume that the start time of the epidemics (introduction of first infected individuals) for the two strains differ by *D*, where *D* is drawn uniformly between [−35,35] days. Once a strain is introduced, it is seeded over a week with 500 cases a day distributed among the different age groups according to the populations sizes and susceptibility to that strain. Thus, 3500 cases are seeded in the population of 318.9 million (estimated US population in 2014).

Once a person is infected with one strain, that person is immune to it for the rest of the outbreak. Moreover, that person’s susceptibility to the other strain is reduced by θ %.

Table 2 summarizes all the concepts introduced in the Materials and Methods. Parameters θ, *V*, *L* are the primary parameters *T_1_* (with parameters vaccine valency *V* and coverage levels *L* used in the comparisons in Table 1), parameters *n_i_*, *c_ij_*, 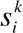, *w*(), λ, *D*, *E* are secondary parameters *T*_2_, as described in the Outline sub-section.

**Table 2:**
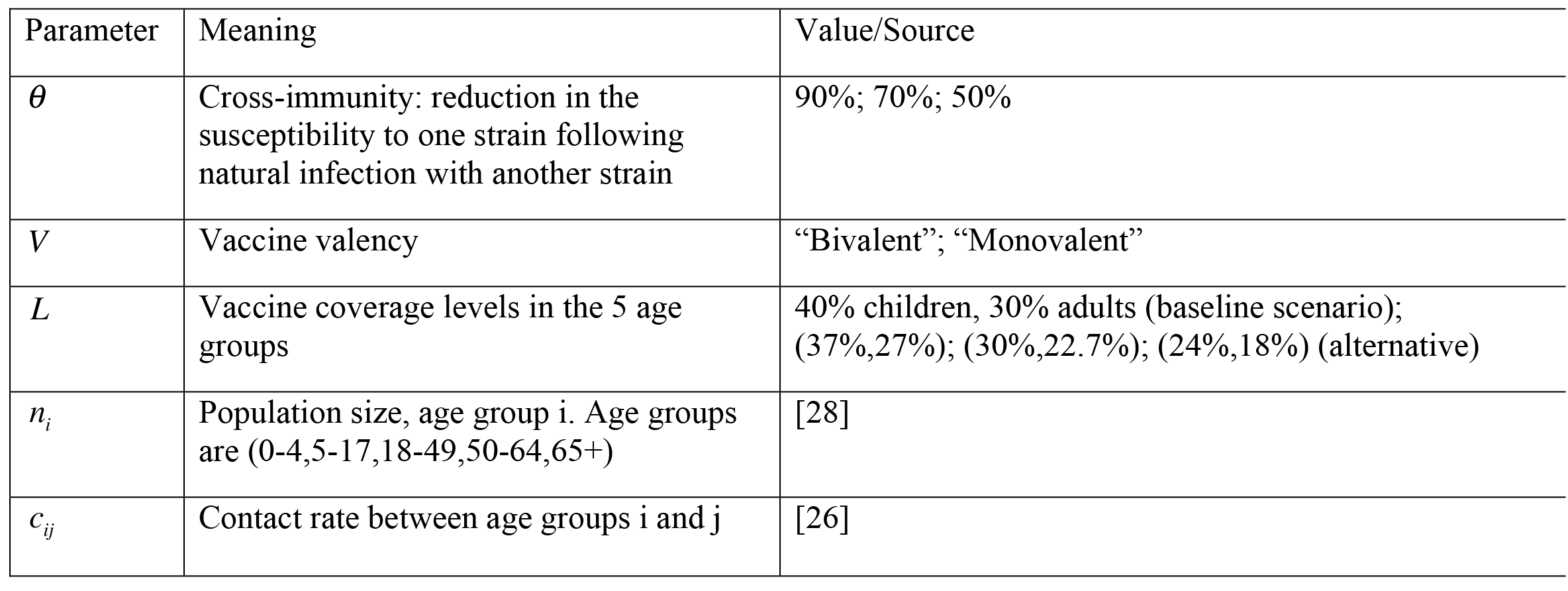
Parameters in the transmission process

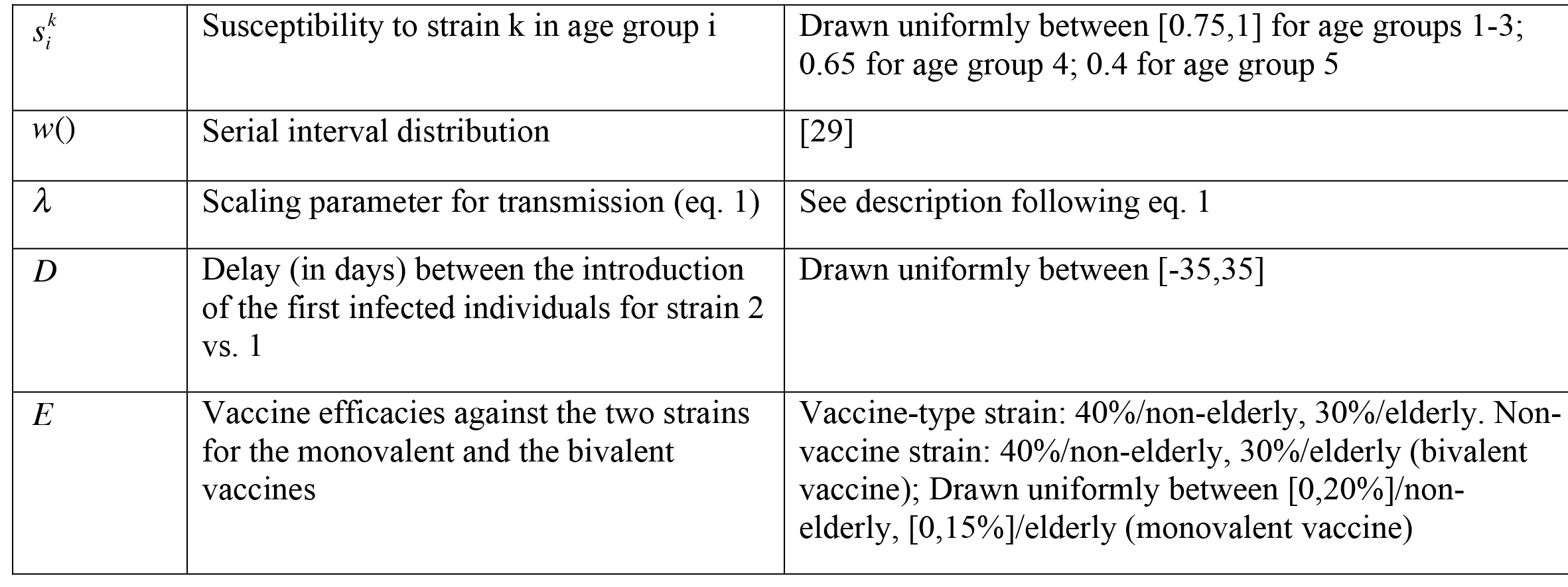

## Results

Figures 1 and 2 give a visual representation of all the comparisons performed in Table 1 (with Figures 1 covering the first 4 rows of Table 1, and Figures 2 covering the last 4 rows). Each panel in either Figures 1 or 2 represents an alternative vaccination policy, and the cumulative incidence under the alternative is plotted against incidence simulated under the baseline coverage of the monovalent vaccine. Thus each point in those plots corresponds to the choice of the other parameters *T*_2_ from the collection *S* (see the Outline sub-section of the Methods) with such a choice defining an epidemic for both the baseline coverage of the monovalent vaccine, and the alternative scenario from Table 1. We also note that points falling under the diagonal represent outbreak scenarios in which the alternative vaccination strategy reduced the size of the epidemic relative to the baseline scenario.

**Figure. 1:**
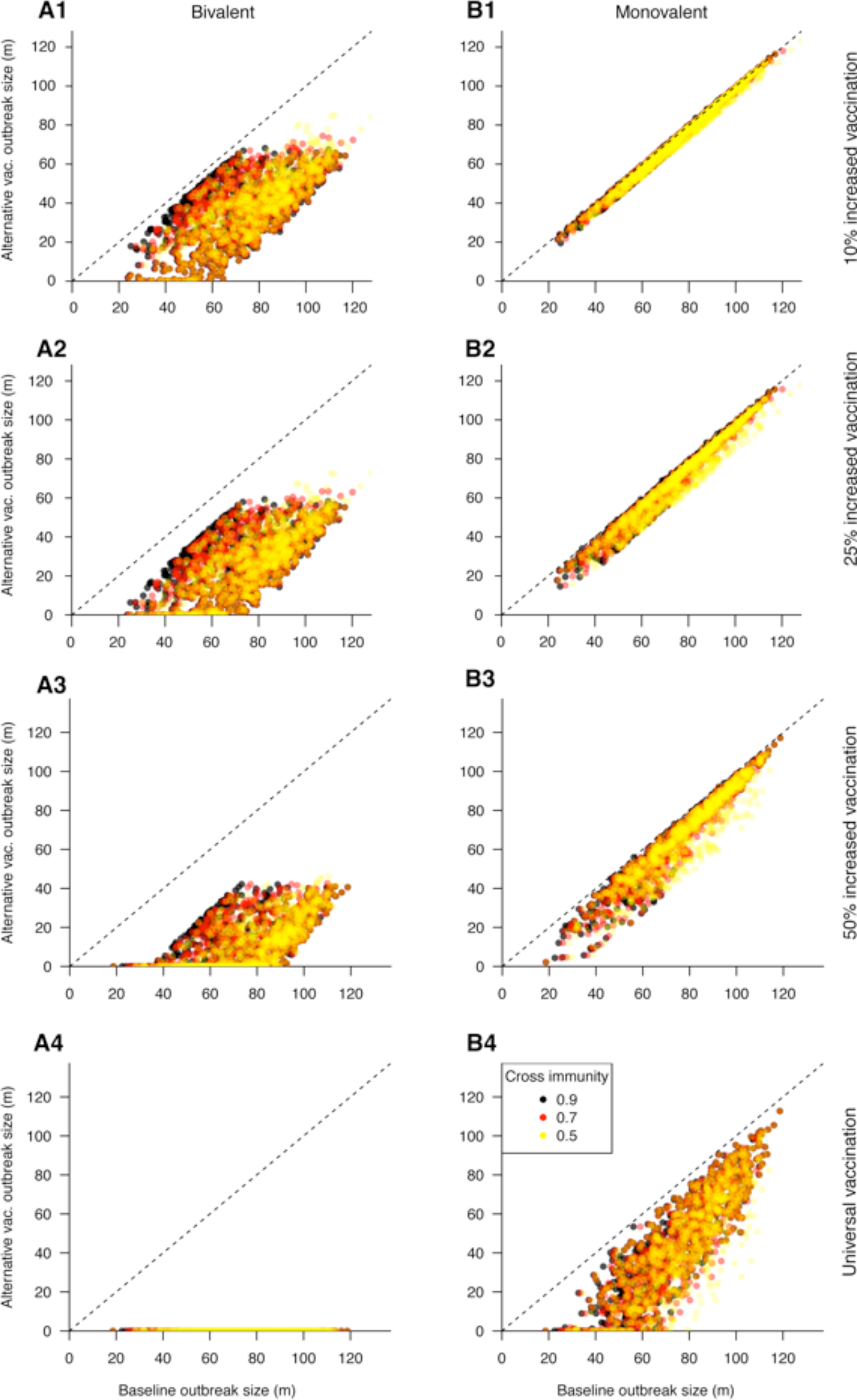
Cumulative incidence (in millions) for simulated epidemics (y-axis) when vaccination coverage is increased (by 10% -Row 1; 25% -Row 2; 50% -Row 3; Universal vaccination -Row 4) vs. the corresponding epidemics for the baseline case.

**Figure. 2.**
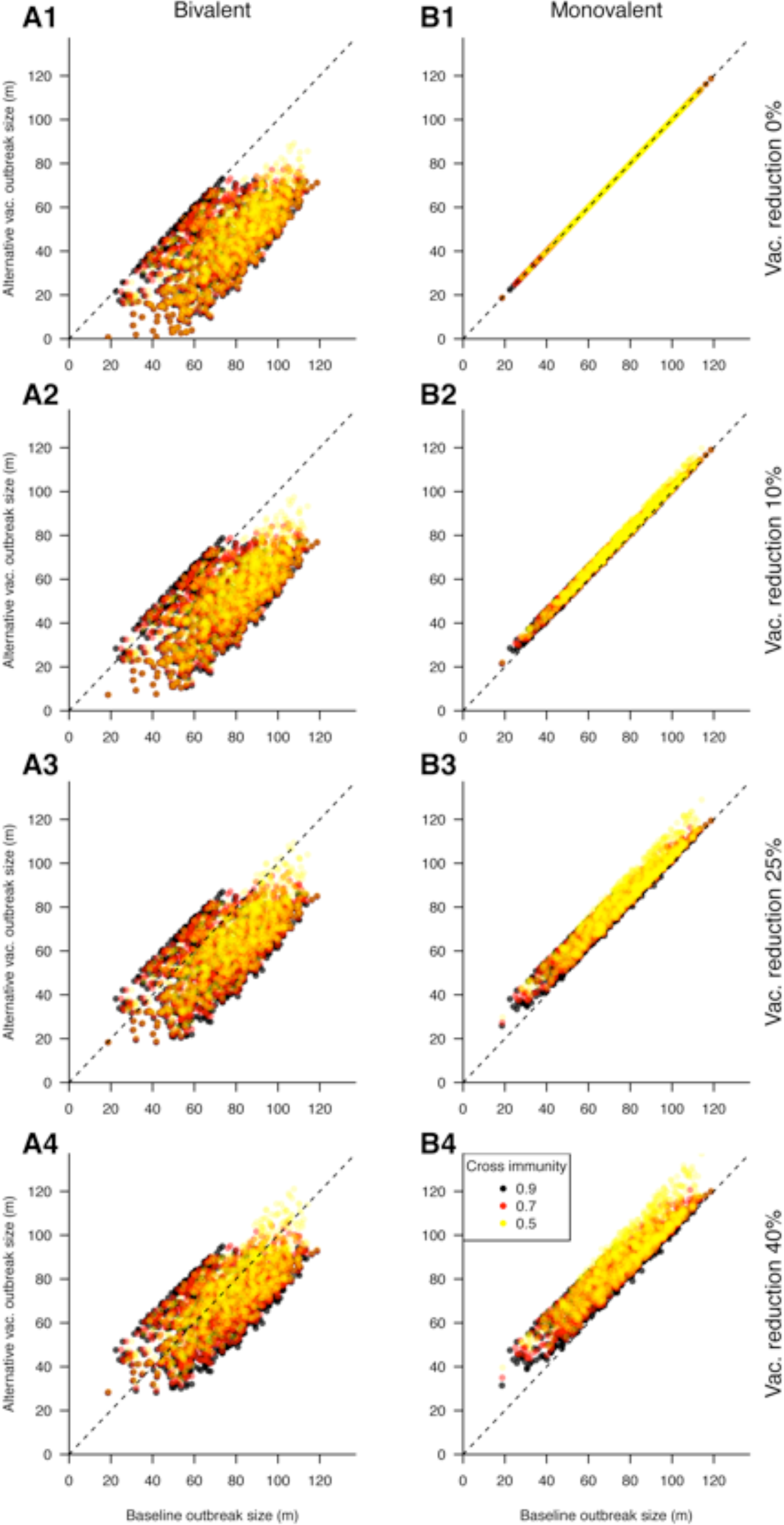
Cumulative incidence (in millions) for simulated epidemics (y-axis) when vaccination coverage is decreased (Unchanged -Row 1; by 10% -Row 2; 25% -Row 3; 40% -Row 4) vs. the corresponding epidemics for the baseline case.

Table 3 compares the modeled epidemiologic outcomes with a bivalent vaccine at varying coverage levels, against that of baseline coverage of a monovalent vaccine. Average cumulative incidence for epidemics with a bivalent vaccine is compared to the baseline case, and the proportion of epidemics with a bivalent vaccine for which the cumulative incidence is smaller than in the baseline scenario (points below the diagonal for the corresponding panels in Figures 1 for the bivalent vaccine) is presented. The results suggest that administration of the bivalent vaccine, even at coverage levels that are 25% lower than the baseline scenario for the monovalent vaccine, leads on average to a significant reduction in incidence compared to the baseline coverage of a monovalent vaccine. Higher coverage levels of the bivalent vaccine prevent the majority of incident cases of infection compared to the baseline coverage of a monovalent vaccine. Moreover, the benefit of the bivalent vaccine increases with increasing strength of cross-immunity.

**Table 3:** Comparative epidemiologic outcomes with the bivalent vaccine coverage from Table 1 compared to the baseline coverage of a monovalent vaccine.

|  | Cross immunity 90% |  | Cross immunity 70% |  | Cross immunity 50% |  |
| --- | --- | --- | --- | --- | --- | --- |
| Coverage for bivalent vaccine relative to baseline | % epidemics with lower cumulative incidence than baseline | Average epidemic size compared to baseline (in %) | % epidemics with lower cumulative incidence than baseline | average epidemic size compared to baseline (in %) | % epidemics with lower cumulative incidence than baseline | average epidemic size compared to baseline (in %) |
| Uniform coverage | 100% | -99.9% | 100% | -99.9% | 100% | -100% |
| 50% increase | 100% | -83.6% | 100% | -83.7% | 100% | -83.9% |
| 25% increase | 100% | -66.5% | 100% | -65.9% | 100% | -65.8% |
| 10% increase | 100% | -51.7% | 100% | -51.2% | 100% | -50.6% |
| Baseline level | 97.1% | -47.2% | 100% | -46.4% | 100% | -45.5% |
| 10% reduction | 88.6% | -37.3% | 93.3% | -36.1% | 96.7% | -34.5% |
| 25% reduction | 82.4% | -23.3% | 84.9% | -21.3% | 88.8% | -18.5% |
| 40% reduction | 73.9% | -10.7% | 71.6% | -7.8% | 61.3% | -3.7% |

Table 4 compares the analogous outcomes for the monovalent vaccine at different coverage levels (Table 1), against the same baseline scenario. The results suggest that cumulative incidence of infection has limited sensitivity to the coverage levels of the monovalent vaccine. Indeed, reductions of a similar magnitude to the administration of a bivalent vaccine at baseline levels could not be attained through increasing monovalent vaccine distribution alone, except the case of unrealistically high (uniform) coverage. Additionally, the higher is the strength of cross-immunity, the less sensitive is the overall incidence to vaccination coverage levels. Overall, the results in Tables 3 and 4 show that the stronger is the degree of cross-immunity, the less beneficial is the increase in coverage levels for the monovalent vaccine, and the more beneficial is the introduction of the bivalent vaccine.

**Table 4:** Comparative epidemiologic outcomes with the monovalent vaccine at higher or lower coverage levels compared to the baseline coverage level (Table 1).

|  | Cross immunity 90% |  | Cross immunity 70% |  | Cross immunity 50% |  |
| --- | --- | --- | --- | --- | --- | --- |
| Vaccine coverage relative to baseline | % epidemics with lower cumulative incidence than baseline | Average epidemic size compared to baseline (in %) | % epidemics with lower cumulative incidence than baseline | average epidemic size compared to baseline (in %) | % epidemics with lower cumulative incidence than baseline | average epidemic size compared to baseline (in %) |
| Uniform coverage | 100% | -38.5% | 100% | -39.8% | 100% | -41.6% |
| 50% increase | 100% | -10.2% | 100% | -11.8% | 100% | -14.2% |
| 25% increase | 100% | -5.6% | 100% | -5.9% | 100% | -6.9% |
| 10% increase | 99.6% | -2.3% | 100% | -2.8% | 100% | -3.5% |
| 10% reduction | 0.8% | +2.5% | 0% | +3.1% | 0% | +4% |
| 25% reduction | 0.9% | +6.4% | 0.1% | +8.1% | 0% | +10.6% |
| 40% reduction | 0.9% | +10.6% | 0.1% | +13.5% | 0% | +17.8% |

In addition to the principal conclusions described in the previous two paragraphs, Tables 3 and 4 suggest that in some (rare) cases, improvements in vaccination policy can be seemingly counterintuitive. Table 3 shows that replacing a monovalent vaccine with a bivalent vaccine at the same coverage level does not always result in a reduction in cumulative incidence. Similarly, Table 4 shows that reducing vaccination coverage for the monovalent vaccine can, under particular circumstances result in larger outbreaks.
The magnitude of those increases in the cumulative incidence of infection can be assessed from Figures 2. Figures 2 (panels A1, B2, B3, B4) suggests that those (rare) increases in the cumulative incidence (points above the diagonal) are quite small, with the magnitude of those increases in cumulative incidence (compared to the baseline) being somewhat larger for the reduction in coverage levels for the monovalent vaccine (panels B2-B4) compared to the introduction of a bivalent vaccine (panel A1).

## Discussion

During some of the past influenza epidemics in the US, co-circulation of a vaccine-type and non-vaccine strains took place. For some of those seasons, incidence of non-vaccine strains increased significantly relative to the incidence of vaccine-types strains; during other seasons, little relative change in the incidence of the different strains was observed.

Administration of vaccines (which often have higher efficacy against vaccine-type strains compared to non-vaccine strains), combined with cross-immunity from natural infections (which is expected to be high) can impact the relative dynamics of the two strains. Vaccine can decrease the incidence of the vaccine-type strain, mitigating its effect on the incidence of the non-vaccine strain.

In this paper, we examine the interaction of cross-immunity and vaccination for the dynamics of co-circulating strains by numerical simulations in an age-stratified population. We consider two types (valencies) of vaccines: bivalent, giving equal (and good) protection against both strains vs. monovalent, with good efficacy against the vaccine-type strain and poor efficacy against the non-vaccine strain. We randomly generate a variety of choices of transmission parameters and consider simulated epidemics with each of those choices of transmission parameters as well as the choices of cross-immunity, vaccination coverage levels and vaccine valency. In our simulations, epidemics in the baseline scenario of a monovalent vaccine (which is meant to reflect the realism of epidemics where vaccine was poor for the non-vaccine strain, e.g. the 2004–05 and 2014–15 A/H3N2 epidemics) and vaccination coverage levels of 40% for children, 30% for adults are matched to epidemics for which vaccine type and coverage levels vary, while all other parameters are the same. This (pair-wise) comparison of the baseline and matched epidemics is used to demonstrate that administration of a bivalent vaccine results in a significant reduction in the overall incidence compared to administration of a monovalent vaccine, oftentimes even when coverage levels of the bivalent vaccine decrease. Moreover, the higher is the degree of cross-immunity, the smaller is the reduction in the incidence of infection that can result from increases in coverage levels for the monovalent vaccine (Table 4), and the more beneficial is the usage of a bivalent vaccine (Table 3). These results are primarily meant to suggest the benefit of including multiple A/H3N2 strain in a vaccine when such strains are expected to circulate.

Our main results are consistent with two simple explanatory principles. The qualitative finding in Table 3 is that when two strains compete for hosts, it is more beneficial to use a bivalent than a monovalent vaccine. Moreover, this benefit generally increases with increasing degree of cross-immunity between the different strains (which is expected to be high, as indicated in the Introduction). We believe that the reason for that is that usage of a monovalent vaccine reduces the incidence of one strain, decreasing the mitigating effect of that incidence on the incidence of the other strain, with the strength of mitigation being highest for higher degree of cross-immunity. Table 4 shows that reducing coverage of the monovalent vaccine typically reduces the population-level benefit of vaccination; however, this reduction of benefit is less striking when cross-immunity is strong, as the increase in incidence of the vaccine-type strain is partly offset by a decline in that of the second strain. We also note that while all those rules hold on average, neither of these rules of thumb holds universally.

This work is meant to illustrate the basic principles underlying the interaction of cross-immunity and vaccination under co-circulation of different strains, rather than make claims about the actual past influenza epidemics in the US. During those epidemics, even the simpler question of which strain would have dominated had the vaccine not been administered is not easy to answer, much less predict in advance. For example, during the process of vaccine selection for the 2014–15 season, the A/Switzerland/9715293/2013 A/H3N2 strain was already known to circulate, but the A/Texas/50/2012 A/H3N2 strain was chosen, with the non-vaccine strain outstripping the vaccine-type strain through the course of the season. In Europe, where vaccination levels are lower than in the US, higher levels of circulation of the vaccine-type A/H3N2 strain took place compared to the US ([31] vs. [6]). We note that regardless of the question which A/H3N2 strain would have been more dominant in the absence of vaccination during the 2014–15 season, and what the impact of the administered vaccine was, it is clear that a vaccine that contained both the Switzerland/2013 and the Texas/2012 A/H3N2 strains would have been significantly more beneficial that a vaccine that only contained one of those strains.

Our work has several limitations. It is unclear how well the range of transmission parameters employed here reflects the reality of influenza epidemics. In our model, vaccine is administered prior to the beginning of influenza seasons while in reality, some additional vaccine administration continues to take place through the course of influenza epidemics, at least in the US. The effect of seasonal forcing on the transmission parameters is not modeled in our study, though this effect should operate independently of the phenomena examined here and presumably has a rather limited impact on the results. While we’ve considered three different values for the strength of cross-immunity parameters, and evidence suggests that this parameter should be fairly large, it is difficult to estimate using epidemiological or genetic data. Moreover, in our simulations this parameter was selected independently of the efficacy of the “monovalent” vaccine against the non-vaccine strain. Given that significant cross-immunity exits between different influenza (sub)types, this assumption for strains within an influenza (sub)type is not unreasonable, though possibly not entirely accurate. Finally, we’ve considered the impact of vaccination on influenza incidence during one season. Incidence of influenza during a given season has an effect on incidence during subsequent seasons through longterm immunity, and correspondingly, vaccination has multi-season effects on incidence as well, even if direct immunity conferred by the vaccine wanes.

## Conclusions

We believe that despite some limitations described in the last paragraph of the Discussion, our work provides a computational framework for understanding several general principles related to vaccination (including the administration of bivalent vaccines) during co-circulation of different influenza strains. It illustrates the significant limitations that monovalent vaccines (those that have poor efficacy against non-vaccine strains) carry and suggests a major improvement in outcomes when bivalent vaccines are administered. We hope that this work can be used to guide vaccine selection. When there is evidence (either based on epidemiological data and/or on the analysis of the evolution of the influenza virus, e.g. [32,33]) that major circulation on multiple, reasonably distinct influenza strains is likely during the upcoming season, our work stresses the benefit of including multiple strains within an influenza (sub)type, especially influenza A/H3N2, in a vaccine.

## Acknowledgement

We thank Jennifer Michaels, Sherida Kipp, and Brian Arnold for their help with this paper.

## References

1. Shaman J, Pitzer VE, Viboud C, Grenfell BT, Lipsitch M (2010) Absolute humidity and the seasonal onset of influenza in the continental United States. PLoS Biol 8: e1000316.

2. Smith DJ, Lapedes AS, de Jong JC, Bestebroer TM, Rimmelzwaan GF, et al. (2004) Mapping the antigenic and genetic evolution of influenza virus. Science 305: 371-376.

3. Ampofo WK, Azziz-Baumgartner E, Bashir U, Cox NJ, Fasce R, et al. (2015)Strengthening the influenza vaccine virus selection and development process: Report of the 3rd WHO Informal Consultation for Improving Influenza Vaccine Virus Selection held at WHO headquarters, Geneva, Switzerland, 1–3 April 2014. Vaccine 33: 4368–4382.

4. US CDC. FluView. 2014–2015 Influenza Season Week 46 ending November 15, 2014.http://www.cdc.gov/flu/weekly/weeklyarchives2014-2015/week46.htm.

5. US CDC. FluView. 2014–2015 Influenza Season Week 50 ending December 13, 2014. http://www.cdc.gov/flu/weekly/weeklyarchives2014-2015/week50.htm.

6. US CDC. FluView. 2014-2015 Influenza Season Week 20 ending May 23, 2015. http://www.cdc.gov/flu/weekly/weeklyarchives2014-2015/week20.htm.

7. US CDC. Seasonal Influenza Vaccine Effectiveness, 2005–2015. http://www.cdc.gov/flu/professionals/vaccination/effectiveness-studies.htm.

8. US Influenza vaccine effectiveness network. End of season influenza vaccine effectiveness estimates for the 2014–15 season. http://www.cdc.gov/vaccines/acip/meetings/downloads/slides-2015-06/flu-02-flannery.pdf.

9. US CDC. Weekly report: Influenza summary update. Week ending January 15, 2005–Week 2. http://www.cdc.gov/flu/weekly/weeklyarchives2004-2005/weekly02.htm.

10. US CDC. Weekly Report: Influenza Summary Update. Week ending May 14, 2005–Week 19 http://www.cdc.gov/flu/weekly/weeklyarchives2004-2005/weekly19.htm.

11. US CDC. Weekly Report: Influenza Summary Update. Week ending December 20, 2003–Week http://www.cdc.gov/flu/weekly/weeklyarchives2003-2004/weekly51.htm.

12. US CDC. Weekly Report: Influenza Summary Update. Week ending January 31, 2004–Week 4 http://www.cdc.gov/flu/weekly/weeklyarchives2003-2004/weekly04.htm.

13. US CDC. Weekly Report: Influenza Summary Update. Week ending May 15, 2004–Week 19 http://www.cdc.gov/flu/weekly/weeklyarchives2003-2004/weekly19.htm.

14. US CDC. Assessment of the effectiveness of the 2003–04 influenza vaccine among children and adults--Colorado, 2003. MMWR Morb Mortal Wkly Rep. 2004 Aug 13;53(31):707–10.

15. US CDC. FluView. 2007–2008 Influenza Season Week 19, ending May 10, 2008. http://www.cdc.gov/flu/weekly/weeklyarchives2007-2008/weekly19.htm.

16. US CDC. FluView. 2007–2008 Influenza Season Week 8, ending February 23, 2008. http://www.cdc.gov/flu/weekly/weeklyarchives2007-2008/weekly08.htm.

17. US CDC. FluView. 2011-2012 Influenza Season Week 20 ending May 19, 2012. http://www.cdc.gov/flu/weekly/weeklyarchives2011-2012/weekly20.htm.

18. US CDC. FluView. 2011-2012 Influenza Season Week 4 ending January 28, 2012. http://www.cdc.gov/flu/weekly/weeklyarchives2011-2012/weekly04.htm.

19. Sonoguchi T, Naito H, Hara M, Takeuchi Y, Fukumi H (1985) Cross-subtype protection in humans during sequential, overlapping, and/or concurrent epidemics caused by H3N2 and H1N1 influenza viruses. J Infect Dis 151: 81–88.

20. Cowling BJ, Ng S, Ma ES, Cheng CK, Wai W, et al. (2010) Protective efficacy of seasonal influenza vaccination against seasonal and pandemic influenza virus infection during 2009 in Hong Kong. Clin Infect Dis 51: 1370–1379.

21. Cowling BJ, Nishiura H (2012) Virus interference and estimates of influenza vaccine effectiveness from test-negative studies. Epidemiology 23: 930–931.

22. Ferguson NM, Galvani AP, Bush RM (2003) Ecological and immunological determinants of influenza evolution. Nature 422: 428–433.

23. Tria F, Lassig M, Pelitl L, Franz S (2005) A minimal stochastic model for influenza evolution. J Stat Mech Theor Exp P07008.

24. Goldstein E, Cobey S, Takahashi S, Miller JC, Lipsitch M (2011) Predicting the epidemic sizes of influenza A/H1N1, A/H3N2, and B: a statistical method. PLoS Med 8: e1001051.

25. Dietz K (1979) Epidemiologic interference of virus populations. J Math Biol 8: 291–300.

26. Mossong J, Hens N, Jit M, Beutels P, Auranen K, et al. (2008) Social contacts and mixing patterns relevant to the spread of infectious diseases. PLoS Med 5: e74.

27. Wallinga J, Teunis P, Kretzschmar M (2006) Using data on social contacts to estimate age-specific transmission parameters for respiratory-spread infectious agents. Am J Epidemiol 164: 936–944.

28. US CDC Wonder. http://wonder.cdc.gov/.

29. Cauchemez S, Donnelly CA, Reed C, Ghani AC, Fraser C, et al. (2009) Household transmission of 2009 pandemic influenza A (H1N1) virus in the United States. N Engl J Med 361: 2619–2627.

30. Worby CJ, Kenyon C, Lynfield R, Lipsitch M, Goldstein E (2015) Examining the role of different age groups, and of vaccination during the 2012 Minnesota pertussis outbreak. Sci Rep 5: 13182.

31. ECDC. Summarising the 2014-2015 influenza season in Europe. http://ecdc.europa.eu/en/press/news/layouts/forms/NewsDispForm.aspx?ID=1231&List=8db7286c-fe2d-476c-9133-18ff4cb1b568-sthash.Duog50bJ.dpuf.

32. Neher RA, Russell CA, Shraiman BI (2014) Predicting evolution from the shape of genealogical trees. Elife 3.

33. R. A. Neher, T. Bedford, R. S. Daniels, C. A. Russell, B. I. Shraiman Prediction, dynamics, and visualization of antigenic phenotypes of seasonal influenza viruses. http://arxiv.org/abs/1510.01195.

